# A water-based platform test of paired or group social dominance and neural activity in male and female mice

**DOI:** 10.1101/2025.07.29.667516

**Authors:** Sarah E Mott, Arushi Saha, Abir Mohsin, Sarah N Magee, Shyenne Grady, Mia N Keller, Melissa A Herman

## Abstract

Social hierarchy is an evolutionarily-conserved phenomenon determined by social dominance behavior that has profound influence on health and relevance in neuropsychiatric disorders. Despite this, the neural mechanisms underlying social dominance remain unclear and current behavioral tests are limited. Here, we describe a novel platform test of social dominance where mice compete for space on a small, elevated platform surrounded by cold water and rank is calculated by total time spent on platform. To validate this assay, we conducted tube test followed by platform test in cages of male and female mice. We observed stable rank in both sexes using the platform test and saw notable differences in overall cage hierarchy between tube and platform. Motivated behavior as measured by attempts to get on the platform was reduced over time. Cooperative behavior as measured by shared time on platform was observed across all ranks but predominantly demonstrated by subordinate mice. Corticosterone levels were significantly higher in females but showed no rank-specific differences. Neuronal activity in the prefrontal cortex was not rank-dependent, however activity in the habenula scaled with rank in both sexes and was significantly correlated with dominance in males. We also investigated group social dynamics using the platform test and identified stable hierarchies with more exaggerated dominance compared to paired platform testing. These results introduce the platform test as a novel method for assessing social dominance behavior in male and female mice, which can be used to further our understanding of the neurobiology underlying social hierarchy.

## Introduction

Social hierarchies form within most animal groups and are characterized by self-organization of individuals into dominant or subordinate ranks^1,2^. Social rank has significant influence on daily behavior, access to resources, and physical and mental health^2,3^. For instance, lower ranks are susceptible to subordination stress, negative health outcomes, shorter lifespans, and increased drug consumption compared to dominant ranks^,5,6,7,8^. Atypical social functioning is a feature of various neuropsychiatric disorders in humans, including anxiety, depression, and substance use disorder^9,10,11^. Stability within social hierarchy can provide a “buffering effect” against neuropsychiatric disease as well as promote reproductive success, conservation of resources, and maximized survival for the entire group^2,3^. However, our current understanding of the neural mechanisms underlying social hierarchy is limited, especially in females. Robust and reliable behavioral tests are essential to investigate the neurobiology of social hierarchy and capture the complexities of social dominance behavior in both sexes.

Various preclinical behavioral assays have been used to measure social hierarchy in rodents^4,12–23^. These assess relative rank through either agonistic encounters or competition for access to resources^12,23^. In agonistic encounters, offensive (dominant) and defensive (subordinate) behaviors are quantified ^12,15,22–24^. While agonistic encounters are effective for single-housed or barrier-housed male mice, they are less reliable at measuring hierarchy in female mice and can involve physical injuries, which may be stressful and impact experimental outcomes^12,22–26^. Resource competition assays are less aggression-based and can determine hierarchy in both sexes^3,12–14,18,21^. In resource-based competition assays, increased access to the resource is equated with greater dominance and is used to determine social rank^1–3,12–14,16,18,21^. Several resource-based competition assays exist, including standard/palatable chow or water competition, the tube test, the wet bedding assay, the warm spot test, and shock avoidance^3,12–14,18,21,27^.

The tube test is the standard resource-based competition assay, which relies on the ethologically relevant resource of space^1–3,12–14,16^. The tube test involves a dominant mouse forcing a subordinate mouse backwards out of a tube and produces clear binary outcomes of social rank in male and female mice^13^. Studies indicate that tube test ranks significantly correlate with home cage dynamics and ranks determined by other dominance metrics^2,12^. Using the tube test, the medial prefrontal cortex (mPFC) and habenula (Hb) have been identified as two brain regions associated with social hierarchy^2,14,27–29^. Activation of mPFC and Hb is associated with dominance^14,27–29^ and manipulation of mPFC activity has been shown to regulate dominance behavior^2,14,27,29^. However, the tube test has limitations resulting from its design that must be considered. Specifically, hierarchy can only be measured between two individuals and multiple pairwise comparisons are necessary to assess group social dynamics^13^, which may not accurately reflect hierarchy in a group setting. The confined nature of the tube test and its binary “win/lose” outcome also limit the behavioral metrics that can be measured.

This study introduces the platform test, a novel behavioral assay for assessing social hierarchy. The platform test leverages the innate drive of mice to avoid cold water, which promotes competition for access to the platform. Social hierarchy was determined by total amount of time spent on the platform and mice were ranked by platform time across testing days. Social ranks determined by platform test were compared to those determined by tube test. Attempts to get on the platform and time spent sharing the platform were assessed to measure motivation and cooperative behavior, respectively. Corticosterone was measured to determine relative stress levels. Neuronal activation was assessed via cFos expression in the PFC and Hb to explore potential rank-dependent differences in region-specific activation. Platform testing was also performed in a group setting and social hierarchy was compared between individual and group testing.

## Methods

### Animals

Male and female adult C57BL/6j mice (n=9/sex, 9-11wks, Jackson Labs) and CRF1-Cre mice (n=4/sex/cage, bred in-house) were group-housed (3/cage/sex) in a humidity– and temperature-controlled vivarium on a 12hr light/dark cycle with ad libitum food and water access. All experimental procedures were approved by UNC Institutional Animal Care and Use Committee.

### Social Dominance Testing

#### Tube Test

Mice were trained to walk through a 30cm tube (males: 30mm I.D., females: 26mm I.D.) 4x/day, 2x/side for two days. Testing was performed as previously described^13^ between cagemates using randomized round-robin pairings. The “loser” was defined as having four paws outside of the tube, while the “winner” remained inside the tube. Testing was video recorded for behavior analysis. Tubes were cleaned between matchups. If both mice attempted to back out of the tube, the trial was ended, and cagemates were returned to their home cage before the matchup was initiated again. Daily social rank was calculated by percentage of wins/total matchups and stability was defined as consistent rank for three consecutive days.

#### Platform Test

The testing arena is comprised of a cylindrical plexiglass chamber (h:16in, w:7.5in) with a beaker (h:2.25in, w:1.5in) filled with colored clay secured in the center surrounded by 1in of cold water (23-25°C). During training, mice were placed in the test chamber and given 1min to find the platform. Paired testing was performed between cage mates in the same randomized round-robin pairings as tube test. Paired testing consisted of two cage mates being placed into the water and given 10min to compete for platform access. Group testing involved all cagemates simultaneously competing for platform access and was performed once daily. Time on platform was defined as having four paws on top of the platform. Platform testing was recorded for subsequent behavior analysis. There were 5min breaks between paired matchups where sides of the test chamber were wiped, and water temperature was measured. Between cages, the chamber was cleaned, and water was replaced. Trials were recorded for behavior analysis. Video recordings of matchups were hand-scored by two investigators. Platform time was averaged by animal across matchups completed each day, yielding an average platform time/day. Social rank was determined by taking average platform time on the final day of testing and sorting it into a designated rank group based on time on platform: 0-200s (subordinate), 201-400s (middle-ranking), or 401-600s (dominant). The number of successful attempts (i.e. getting onto the platform), unsuccessful attempts (i.e. failing to get on the platform), and shared platform time (i.e. time when both subjects were on the platform) were manually scored. Stability was defined as consistent rank for three consecutive days. Paired and group platform testing were analyzed separately, and social ranks were compared.

### Corticosterone Assay

Blood was centrifuged, and serum was collected and stored at −20 °C. Serum was analyzed for corticosterone using a commercially available ELISA kit (Arbor Assays, Ann Arbor, MI, CAT#K014-H5) per the manufacturer’s protocol. Results are expressed as nanograms/milliliter (ng/mL).

### Immunohistochemistry

Mice were anesthetized with isoflurane 90min following testing and perfused with 1x phosphate-buffered saline (PBS) followed by 4% paraformaldehyde (PFA). Brains were postfixed in 4% PFA then transferred to 30% sucrose in PBS and stored at 4°C. Brains were sectioned into 40μm slices using a microtome (HM450, Thermo Fisher Scientific) and stored in 0.01% sodium azide/PBS at 4°C. Slices were washed 2×10min in PBS, incubated for 30min in 50% methanol/PBS, washed 1×5mins in 3% hydrogen peroxide, and incubated for 1hr in blocking solution (0.3% Triton X-100/PBS) and 1% Bovine Serum Albumin. Slices were then incubated for 48hr in rabbit anti-cFos (1:3000, Millipore Sigma; ABE457) at 4°C. After 48hr, slices were washed 1×10min with Tris, NaCl, Triton X-100 (TNT) buffer followed by a 30min wash in Tris, NaCl, and blocking reagent (TNB; PerkinElmer). Slices were incubated for 30min in Horse Radish Peroxide (HRP; 1:200, Abcam ab6721), and washed 4×5min in TNT. Slices were then incubated for 10min in Cy3 [1:50] TSA amplification diluent (Akoya Biosciences, NEL744001KT) at room temperature before being washed 2×10min in TNT. Slices were mounted onto slides using Vectashield DAPI with hard-set (Vector labs; H1500-10). Images were taken on a Keyence BZ-X800 microscope at 20x and the BZ-X800 Analyzer program was used to create stitched images. The PFC and Hb were outlined in ImageJ and quantification was averaged between two blinded investigators.

### Statistical analysis

Statistical analysis and graph generation were performed using Prism 10.0 (GraphPad). Data were analyzed and compared using 1-way ANOVA or a 2-way ANOVA with Tukey’s multiple comparisons or a linear regression. p < 0.05 indicated statistical significance. All data sets are expressed as mean ± SEM.

## Results

The platform test arena is comprised of a cylindrical chamber with one inch of cold water (23-25°C) surrounding a small beaker packed with colored clay (**Figure 1A**). Mice were trained for two days to find the platform before five days of testing. Platform testing involved cagemates competing for platform access for 10min/day/5days (**Figure 1B**). Platform time was measured to determine relative cage hierarchy. Matchups were performed in randomized round-robin pairings within each cage (**Figure 1C**).

**Figure 1.**
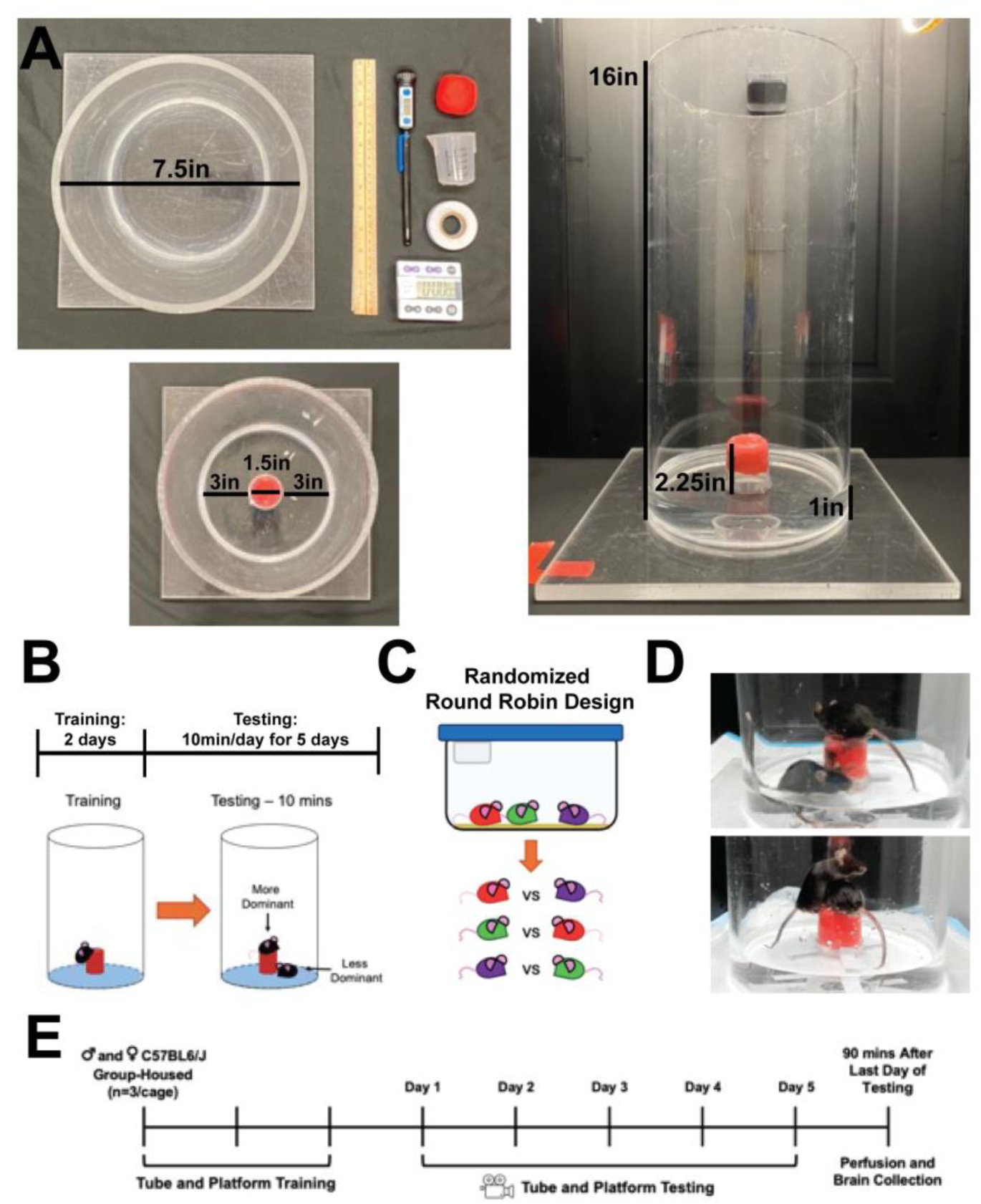
Platform Test Setup and Study Timeline. **A.** The test chamber was constructed with a cylindrical chamber, a beaker, tape, cold water (23-25°C), and colored clay. **B.** Basic timeline of the platform test. **C.** Randomized round robin design used for pairwise matchups. **D.** Dominance behavior (top) and sharing behavior (bottom) were seen on the platform. **E.** Timeline of the study.

Cooperative and motivated behavior were measured through shared platform time and platform attempts, respectively (**Figure 1D**). To compare hierarchies by test, we subjected cages of male and female mice (n=3/cage) to two days of tube and platform training, followed by five days of tube and platform testing (**Figure 1E**).

Using the tube test, social rank was determined for male mice (**Figure 2A**). All cages had clear linear hierarchies with general stability in rank (**Figure 2A, left**). The platform test was then used to assess social rank in the same mice with time on platform used as the metric of dominance (**Figure 2A, right**). All cages had relative stability in rank with each mouse spending roughly the same amount of time on the platform across days (**Figure 2A**). Social hierarchy was more complex, however, with mice displaying proximal ranks as assessed by similar amounts of time on the platform in addition to clear dominant or subordinate ranks (**Figure 2A, right**). When social rank by tube test was measured in females, linear hierarchies were also seen, although hierarchy stability was relatively low with high variability across days (**Figure 2B, right**). The platform test was then performed in the same female mice and high rank stability was seen throughout the testing (**Figure 2B, right**). Female mice had complex hierarchy dynamics in platform testing with most mice displaying proximal ranks (**Figure 2B, right**). Body weights were not associated with tube rank or platform time in either sex (**Figure S1**). Tube ranks were relatively stable in males but showed less stability in females (**Figure 2C**). However, individual platform ranks showed similar stability in both sexes (**Figure 2D**). Female mice were evenly distributed across social rank groups, but male mice were more represented in the middle-ranks (**Figure 2E**). There was a significant main effect of tube rank on platform time in males (**Figure 2F**; 1-way ANOVA: F(2,6)=9.638, p=0.0134) with middle-ranking males spending significantly more time on the platform than subordinate males (1-way ANOVA: Tukey’s: p=0.0114). In females, there was no effect of tube rank on platform time (**Figure 2G**).

**Figure 2.**
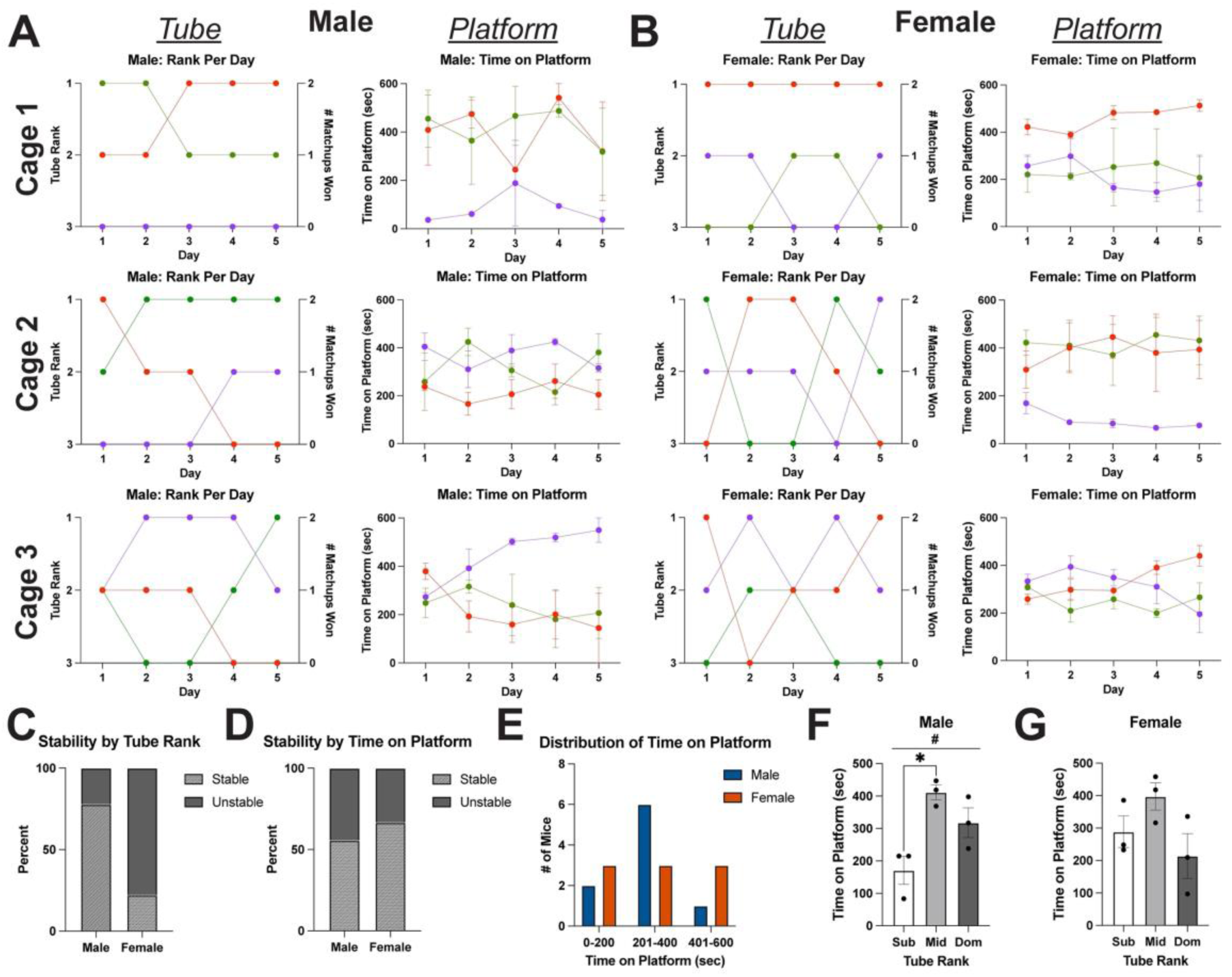
Social Dominance Behavior and Rank Stability. **A.** Hierarchy in 3 cages of male mice (n=9, 3/cage) was assessed using tube test and platform test. **B.** Hierarchy in three cages of female mice (n=9, 3/cage) was measured using tube and platform test. **C.** Stability of individual tube ranks across sex. **D.** Stability of individual time of platform across sex. **E.** Distribution of male and female mice across social rank groups based on time on platform. **F.** Time males spent on the platform based on their tube rank (1-way ANOVA: main effect of rank, F(2,6)=9.638, #p=0.0134; Tukey’s, sub v mid, p=0.0114). **G.** Time females spent on the platform based on their tube rank.

Successful and unsuccessful attempts to get on the platform were quantified during platform testing for male and female mice (**Figure 3A**). On Day 1, there was a significant main effect of platform time on attempts made by males (**Figure 3B**; 2-Way ANOVA: F(2,12)=13.28, p=0.0009) and subordinate males made more unsuccessful attempts than middle-ranking males (**Figure 3B**; 2-way ANOVA: Tukey’s: p=0.0195). Subordinate males also had the highest total number of attempts on Day 1, with significantly more attempts than middle-ranking males (**Figure 3C**, 2-way ANOVA: Tukey’s: p=0.0018). There was a significant interaction between total attempts made by males and day of testing, where attempts were reduced by Day 5 in a rank-dependent manner (**Figure 3C**; 2-way ANOVA: main effect of day: F(1,12)=113.9, p<0.0001; main effect of platform time: F(2,12)=5.489, p=0.0203; interaction: F(2,12)=9.211, p=0.0038). However, the type of attempt made by males was not significantly different on Day 5 (**Figure S2**). In females, there was a significant main effect of platform time on Day 1 attempts (**Figure 3D**, 2-Way ANOVA: F(2,12)=4.889, p=0.0280). Subordinate females had the highest total number of attempts on Day 1, with significantly more attempts than middle-ranking females (**Figure 3E**, 2-way ANOVA: Tukey’s: p=0.0053) and dominant females (2-way ANOVA: Tukey’s: p=0.0053). The type of attempt made by females was not significantly different on Day 5 (**Figure S2**), but total attempts were significantly reduced in females over time in a rank dependent manner with a significant interaction between attempts and day of testing (**Figure 3E**, 2-way ANOVA: main effect of day: F(1,12)=117.7, p<0.0001; main effect of time on platform: F(2,12)=9.144, p=0.0039; interaction: F(2,12)=5.814, p=0.0172). There were no sex differences in total attempts for Day 1 or Day 5 (**Figure S2**).

**Figure 3.**
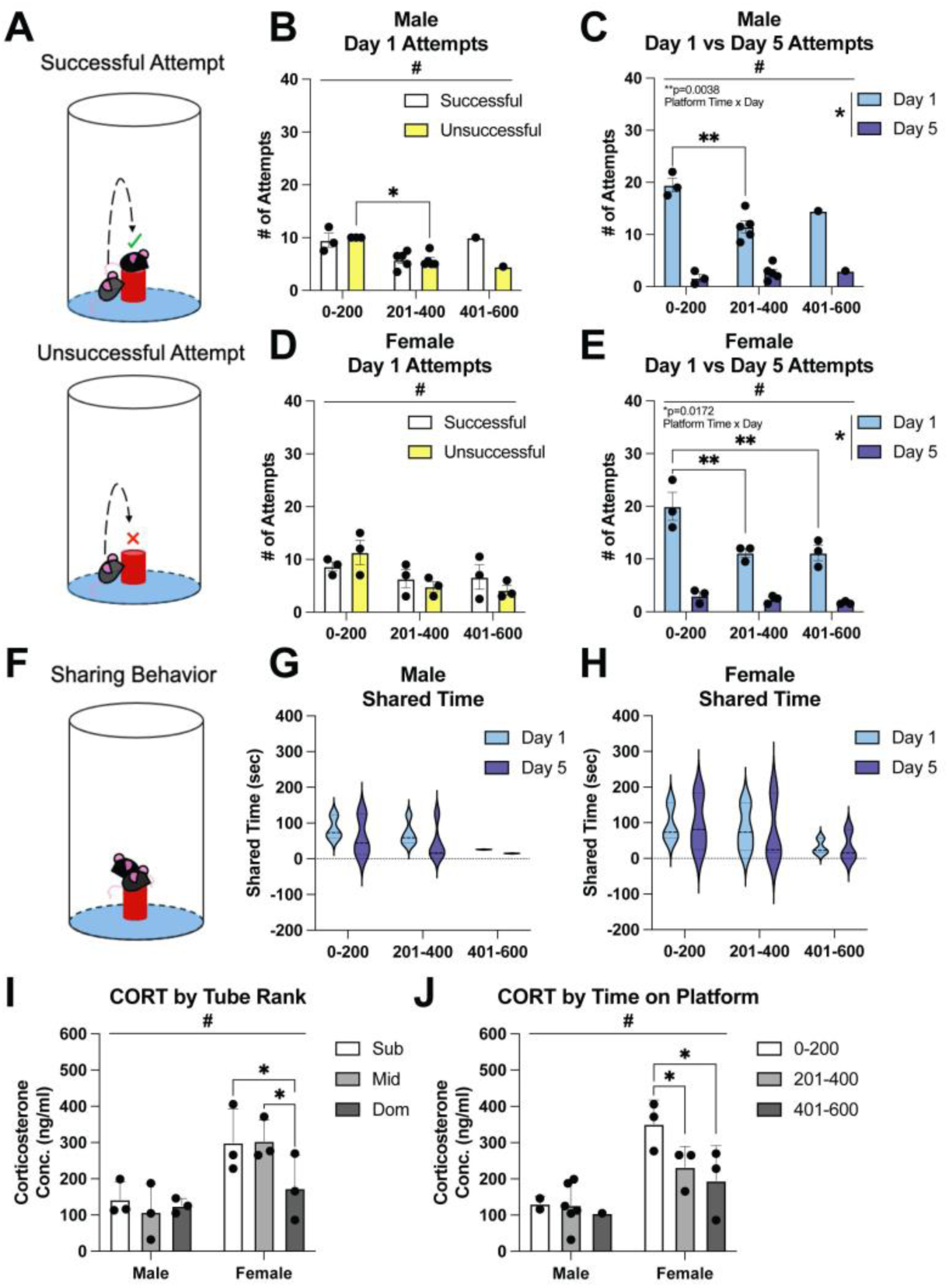
Assessment of Motivation, Cooperative Behavior, and Stress. **A.** Attempts to get onto the platform were categorized as successful or unsuccessful. **B.** Types of attempts made by male mice to get on the platform on Day 1 (2-way ANOVA: main effect of time on platform, F(2, 12)=13.28, #p=0.0009; Tukey’s, 0-200 v 201-400, p=0.0195). **C.** Total attempts made by male mice on Day 1 compared to Day 5 (2-way ANOVA: main effect of day: F(1,12)=113.9, *p<0.0001; main effect of time on platform: F(2,12)=5.489, #p=0.0203; interaction: F(2,12)=9.211, p=0.0038; Tukey’s, 0-200 v 201-400, p=0.0018). **D.** Successful and unsuccessful attempts made by female mice to get on the platform on Day 1 (2-Way ANOVA: main effect of platform time, F(2,12)=4.889, #p=0.0280). **E.** Total attempts made by female mice on Day 1 compared to Day 5 (2-way ANOVA: main effect of day: F(1,12)=117.7, *p<0.0001; main effect of time on platform: F(2,12)=9.144, #p=0.0039; interaction: F(2,12)=5.814, p=0.0172; Tukey’s, 0-200 v 201-400, p=0.0053; Tukey’s, 0-200 v 401-600, p=0.0053). **F.** Cooperative behavior was assessed by time spent sharing the platform. **G.** Total time spent sharing the platform on Day 1 compared to Day 5 in male mice. **H.** Total time female mice spent sharing the platform on Day 1 compared to Day 5. **I.** Corticosterone levels assessed by tube test rank in male and female mice (2-Way ANOVA: main effect of sex, F(1,12)=16.45, #p=0.0016; Tukey’s, Mid v Dom, p=0.0415; Sub v Dom, p=0.0477). **J.** Corticosterone levels assessed by time on platform in male and female mice (2-Way ANOVA: main effect of sex, F(1,12)=14.61, #p=0.0024; Tukey’s, Sub v Mid, p=0.0490; Tukey’s, Sub v Dom, p=0.0138).

Shared time on the platform was measured to assess cooperative behavior (**Figure 3F**). Male mice of all ranks shared the platform, although dominant mice had lower total shared times (**Figure 3G**). In females, sharing was also seen across all ranks with reduced shared time amongst dominant females (**Figure 3H**). There were no sex differences in sharing (**Figure S2**). Male and female corticosterone levels were not significantly altered by tube rank, but there was a significant main effect of sex (**Figure 3I**; 2-Way ANOVA: F(1,12)=16.45, p=0.0016), and dominant females displayed significantly lower corticosterone compared to middle ranking (**Figure 3I**; 2-Way ANOVA: Tukey’s: p=0.0415) and subordinate females (2-Way ANOVA: Tukey’s: p=0.0477). Platform time did not impact corticosterone levels, but there was a significant main effect of sex (**Figure 3J**; 2-Way ANOVA: F(1,12)=14.61, p=0.0024) and subordinate females had significantly higher corticosterone levels compared to middle-ranking (**Figure 3J**; 2-Way ANOVA: Tukey’s: p=0.0490) and dominant females (2-Way ANOVA: Tukey’s: p=0.0138).

Neuronal activity following tube and platform testing was measured via cFos expression in PFC, including anterior cingulate (Cg1 and Cg2), infralimbic (IL), prefrontal (PrL), and medial orbital (MO) subregions to examine any rank-dependent differences (**Figure 4A**). Overall PFC activity did not differ by tube rank or platform time for either sex (**Figure S3**). A significant main effect of PFC subregion on cFos expression was seen in males (**Figure 4B**; 2-Way ANOVA: F(4,28)=6.959, p=0.0005) and in females (**Figure 4C**; 2-Way ANOVA: F(4,26)=9.034, p=0.0001), but no significant differences were seen between tube ranks for either sex. Similarly, neuronal activity did not differ by platform time in either sex, but a significant main effect of PFC subregion was seen in males (**Figure 4D**; 2-Way ANOVA: F(4,28)=4.262, p=0.0081) and in females (**Figure 4E**; 2-Way ANOVA: F(4,26)=8.678, p=0.0001). Neuronal activity in the medial (MHb) and lateral (LHb) Hb subregions was examined for rank-dependent differences (**Figure 4F**). Overall Hb activity did not differ by tube rank or platform time for either sex (**Figure S3**). In males, there was a significant interaction of Hb subregion and tube rank (**Figure 4G**; 2-Way ANOVA: F(2,12)=4.764, p=0.0300) with an overall effect of Hb subregion (2-Way ANOVA: F(1,12)=7.108, p=0.0206), such that dominant mice had significantly higher activity than subordinate mice in LHb (**Figure 4G**; 2-Way ANOVA: Tukey’s, p=0.0496). In females, tube rank did not alter neuronal activity (**Figure 4H**), but there was a significant main effect of Hb subregion (2-Way ANOVA: F(1,12)=12.47, p=0.0041). Neuronal activity was significantly different by platform time in males (**Figure 4I**; 2-Way ANOVA: F(2,12)=6.102, p=0.0149) with a significant interaction of Hb subregion and platform time (2-Way ANOVA: F(2,12)=8.834, p=0.0044). There were no differences in cFos expression by Hb subregion in males (**Figure 4I**), but dominant mice displayed significantly greater activation in LHb (2-Way ANOVA: Tukey’s, p=0.0218). In females, there was a significant interaction of Hb subregion and platform time (**Figure 4J**; 2-Way ANOVA: F(2,12)=5.230, p=0.0233) with an overall effect of Hb subregion (2-Way ANOVA: F(1,12)=10.01, p=0.0082) and platform time (2-Way ANOVA: F(2,12)=5.727, p=0.0179), such that dominant females had greater activation than middle-ranking females in LHb (**Figure 4J**; 2-Way ANOVA: Tukey’s, p=0.0057). Regression analysis of neuronal activation in LHb and platform time found a significant positive association in males (**Figure 4K**; Simple Linear Regression: R^2^=0.4530, p=0.0469).

**Figure 4.**
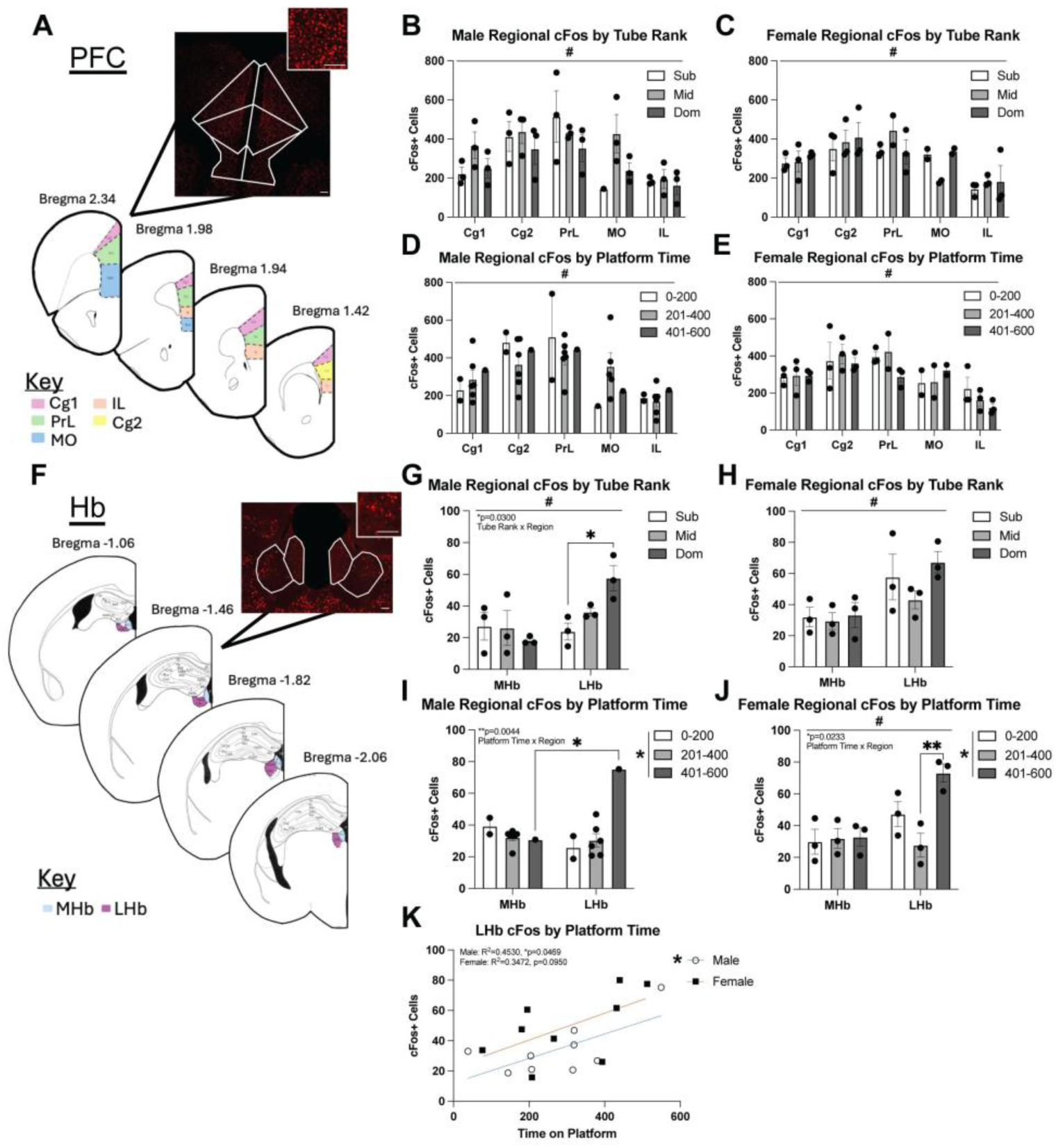
Neuronal Activity Following Tube and Platform Testing in PFC and Hb. **A.** cFos, a marker of neuronal activity, was assessed in subregions of the PFC (Cg1 and Cg2 = anterior cingulate cortex, IL= infralimbic cortex, PrL = prefrontal cortex, and MO = medial orbital prefrontal cortex) following behavioral testing, scale bar = 50um. **B.** Male PFC subregion cFos expression averaged by animal for tube rankings (2-Way ANOVA: main effect of subregion: F(4,28)=6.959, #p=0.0005). **C.** Female PFC subregion cFos expression averaged by animal for tube rankings (2-Way ANOVA: main effect of subregion: F(4,26)=9.034, #p=0.0001). **D.** Male PFC subregion cFos expression averaged by animal for time on platform (2-Way ANOVA: main effect of subregion: F(4,28)=4.262, #p=0.0081). **E.** Female PFC subregion cFos expression averaged by animal for time on platform (2-Way ANOVA: main effect of subregion: F(4,26)=8.678, #p=0.0001). **F.** Neuronal activity in the medial (MHb) and lateral (LHb) subregions of the Hb was quantified following tube and platform testing, scale bar = 50um. **G.** Male Hb subregion cFos expression averaged by animal for tube rankings (2-Way ANOVA: main effect of subregion: F(1,12)=7.108, #p=0.0206; significant interaction: F(2,12)=4.764, p=0.0300; Tukey’s, Sub v Dom, p=0.0496). **H.** Female Hb cFos expression averaged by animal for tube rankings (2-Way ANOVA: main effect of subregion: F(1,12)=12.47, #p=0.0041). **I.** Male Hb subregion cFos expression averaged by animal for time on platform (2-Way ANOVA: main effect of platform time: F(2,12)=6.102, *p=0.0149; significant interaction: F(2,12)=8.834, p=0.0044; Tukey’s, MHb v LHb, p=0.0218). **J.** Female Hb subregion cFos expression averaged by animal for time on platform (2-Way ANOVA: main effect of subregion: F(1,12)=10.01, #p=0.0082; main effect of platform time: F(2,12)=5.727, *p=0.0179; significant interaction: F(2,12)=5.230, p=0.0233; Tukey’s, Mid v Dom, p=0.0057). **K.** Regression analysis of cFos expression in the LHb subregion and platform time for males and females (Simple linear regression: male: R^2^=0.4530, p=0.0469; female: R^2^=0.3472, p=0.0950).

Platform testing was also performed in a group setting, where male and female mice (n=4/cage) were subjected to tube test followed by paired or group platform test (**Figure 5A**). Paired testing involved competition between two cage mates (**Figure 5B**) and group testing involved competition between all cagemates simultaneously (**Figure 5C**). In males, a relatively stable linear hierarchy was seen throughout tube test with a consistent dominant mouse (**Figure 5D, left**). Paired platform testing was also relatively stable, although proximal ranks separated into a more linear hierarchy across days (**Figure 5D, right**). In group platform testing, the dominant male remained, while all other males showed increasingly subordinate positions (**Figure 5D**). Females showed early variability in tube rank, but stable linear hierarchy was identified by Day 9 (**Figure 5E, left**). Paired platform testing maintained stability across days with one subordinate female and proximal dominant ranks amongst other cagemates (**Figure 5E, right**). In group platform testing, dominance was stratified amongst two more dominant females and two proximal subordinate females (**Figure 5E, right**). Body weights were not associated with tube rank or platform time in either sex (**Figure S4**). In males, the type of attempt made on Day 1 was not significantly different and did not depend on platform time (**Figure 5F**). Total attempts by males also did not differ across platform time or across days of paired platform testing (**Figure 5G**). In females, there was a significant main effect of attempt type on Day 1 (**Figure 5H**; 2-Way ANOVA: F(1,4)=65.56, p=0.0013) with a higher number of successful attempts in subordinate (2-Way ANOVA: Tukey’s, p=0.0198) and dominant females (2-Way ANOVA: Tukey’s, p=0.0079). Total attempts made by females were significantly different by platform time (**Figure 5I**, 2-way ANOVA: F(1,4)=10.04, p=0.0339) with a significant interaction of platform time and day of paired platform testing (2-way ANOVA: F(1,4)=13.44, p=0.0215), such that dominant females made more attempts than subordinate females on Day 4 (2-Way ANOVA: Tukey’s, p=0.0283). Moreover, both males and females made a greater number of successful attempts on Day 4, and this was predominantly seen amongst more dominant mice (**Figure S5)**. Time spent sharing did not differ across platform time or across days of paired platform testing in males or females (**Figure S5**). However, sex differences were observed on Day 1 with females making more attempts and sharing more often than males (**Figure S5**). On Day 5, the type of attempt made by males (**Figure 5J**) and females (**Figure 5L)** was not significantly different and did not depend on platform time. Total attempts made by males (**Figure 5K**) and females (**Figure 5M)** also were not dependent on platform time and did not significantly change across days of group platform testing. However, sex differences were observed on Day 5 with females making more attempts than males (**Figure S6**). Time spent sharing was not different across platform time or across group platform testing days (**Figure S6**).

**Figure 5.**
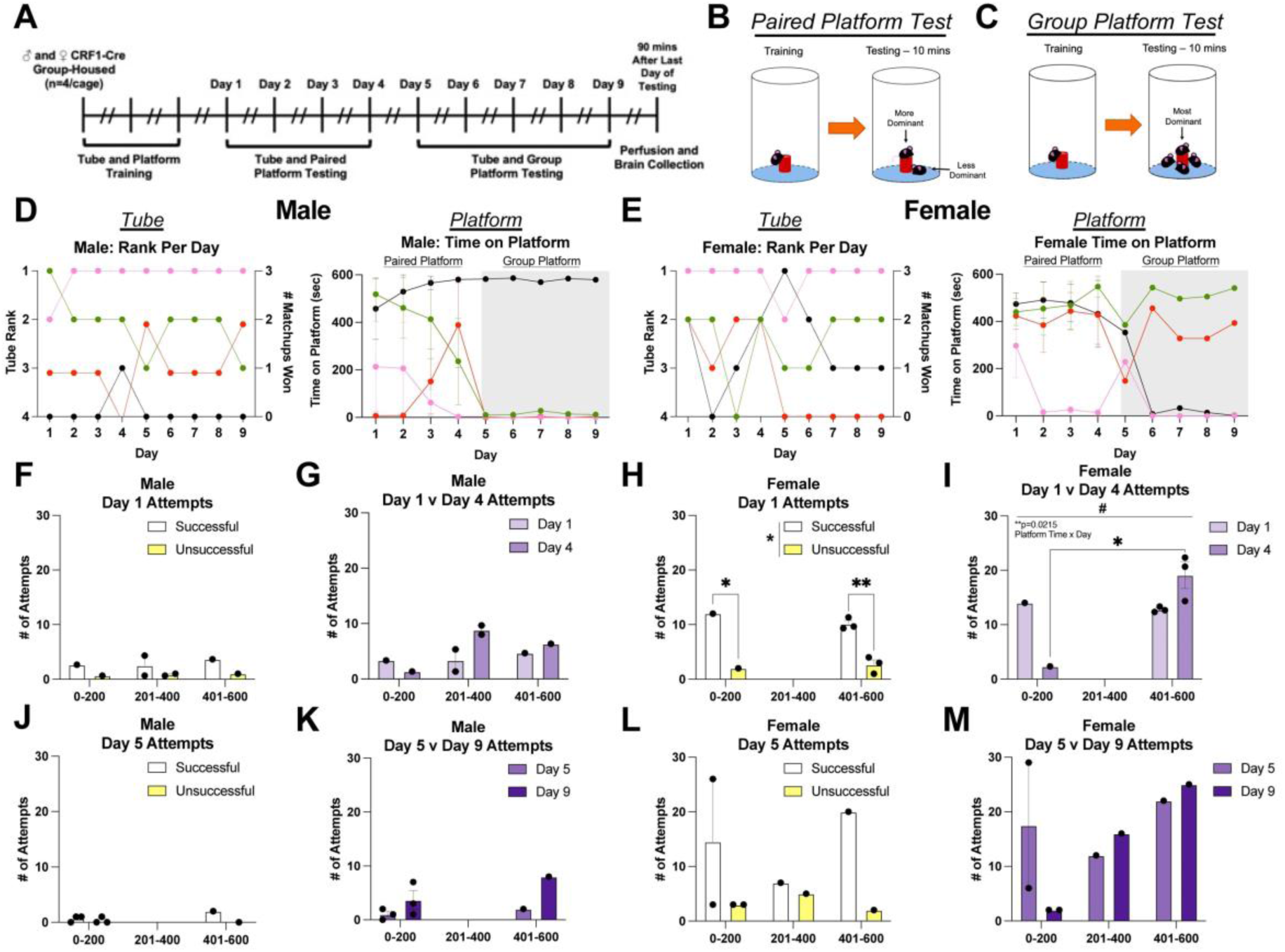
Group Dynamics Measured with Platform Test. **A.** Experimental timeline with 1-3 days between testing. **B.** Paired platform testing was conducted between two cagemates. **C.** Group platform testing was conducted between all cagemates simultaneously. **D.** Male cage dynamics (n=4, 4/cage) measured with tube test, paired platform test, and group platform test. **E.** Female cage dynamics (n=4, 4/cage) measured with tube test, paired platform test, and group platform test. **F.** Types of attempts made by male mice to get on the platform on Day 1. **G.** Total attempts made by male mice across paired platform testing. **H.** Types of attempts made by female mice to get on the platform on Day 1 (2-way ANOVA: main effect of attempt type: F(1,4)=65.56, *p=0.0013; Tukey’s, 0-200 Successful v Unsuccessful, p=0.0198; Tukey’s, 401-600 Successful v Unsuccessful, p=0.0079). **I.** Total attempts made by female mice across paired platform testing (2-Way ANOVA: main effect of platform time: F(1,4)=10.04, #p=0.0339; significant interaction: F(1,4)=13.44, p=0.0215; Tukey’s, 0-200 v 401-600, p=0.0283). **J.** Types of attempts made by male mice to get on the platform on Day 5. **K.** Total attempts made by male mice across group platform testing. **L.** Types of attempts made by female mice to get on the platform on Day 5. **M.** Total attempts made by female mice across group platform testing.

## Discussion

The goal of this study was to develop the platform test, an alternative resource-based competition assay to investigate social hierarchy in male and female mice. In the platform test, time on platform determines relative social rank and motivated and cooperative behavior are assessed. The platform test produced reliable social ranks in both sexes and found hierarchy complexity with proximal ranks that were not identified using tube test. The platform test showed stable hierarchies for both sexes, while tube test was only stable amongst males. Attempts made by male and female mice were dependent on platform time and were reduced across days of testing. Analysis of cooperative behavior showed that both sexes shared the platform, but dominant mice spent less time sharing. PFC and Hb activity following behavioral testing was measured by cFos expression. PFC activity was significantly different by subregion but not rank. Hb activity was dependent on time on platform for both sexes, but only dependent on Hb subregion in females. Regression analysis indicated a significant association of activation in the LHb and platform time in males. Group dynamics were also measured in both sexes. There was more extreme stratification of the hierarchy across paired and group testing, but consistency in dominant ranks. Together, this work demonstrates that platform test can be used to assess hierarchy in mice of both sexes in a paired or group setting with additional metrics of behavioral analysis.

The tube test is a standard assay for measuring social hierarchy in mice^1–3,12–14,16^. In this study, tube test identified linear hierarchies, but platform test identified a more complex spectrum of social dynamics. Comparison of social ranks showed notable differences in hierarchy between tests. These differences may be due to the order of testing as tube test was consistently performed prior to platform test. They may also be the result of different test parameters or behavioral metrics; tube test is a brief head-to-head confrontation with a binary “win/lose” outcome^13^ and platform test is a prolonged dominance interaction with rank determined by platform time. It is possible that the tube test was artificially imposing a linear hierarchy due to its binary metrics, while platform test captured a wider spectrum of dominance. This spectrum of dominance may be particularly important in female mouse hierarchy, which is understudied and shows conflicting linearity in the literature^1,30,31^. As strain-specific and rodent-specific differences exist in social behavior and hierarchy, future work should explore the ability of the platform test to measure social dominance behavior in other rodent species and strains^33^. Our work found that female platform test showed stability in platform time, while tube ranks were highly variable. This was not a result of our five-day testing window as both sexes have been shown to establish stable hierarchies rapidly^32^. Moreover, it is unlikely that platform hierarchy is the result of learning specific to the test itself as platform time remained consistent across days in both sexes, indicating the assay is effective at assessing stable hierarchies.

The platform test assesses motivation and cooperation, which cannot be measured with tube test due to its binary outcomes. Recent tube test work has expanded test metrics to include behaviors in the tube, like pushing, resistance, and retreat^13,27,29^. However, it remains unclear whether these additional tube measurements effectively gauge the internal state of the mouse^14,34^. The platform test leverages the innate drive of mice to avoid cold water, which allows for assessment of motivation by attempts to get on the platform. We saw no significant differences in successful or unsuccessful attempts in either sex, but total attempts were reduced across testing days. While food or water competition assays have quantified successful attempts as a metric of rank^12^, this is the first test using both successful and unsuccessful attempts to better understand relative motivation to maintain the resource. We also found that both sexes cooperatively shared the platform, but there was more sharing in females and reduced cooperation in dominant mice. This aligns with studies showing that mice exhibit cooperative behavior in a social setting and that dominance can regulate cooperation by establishing power dynamics that coordinate actions within a group^35,36^. The platform test was also able to measure group dynamics in mice, which few existing assays can do. We found stable social ranks with group platform testing, which paralleled our findings in paired matchups, although hierarchy differences were magnified. The hierarchy differences between paired and group matchups are significant because pairwise social dynamics do not necessarily reflect social dominance that occurs in a larger group context. Moreover, platform test can be used to assess the relative role of different brain regions or circuits in dominance behavior. We found that LHb, but not PFC activation was associated with rank in both tube and platform. This supports previous literature implicating the LHb^28^ but opposes work associating PFC activation with dominance^2,14,27–29^. This may be due to differences in study timelines or rodent strains.

Social hierarchy significantly impacts health and disease^2,3,9–11^, making it critical that robust and reliable behavioral tests exist to study underlying neurobiology. Our work introduces the platform test to measure social hierarchy in male and female mice. We find that the platform test produces stable hierarchies in both sexes, provides additional depth of behavioral analysis, and assesses paired or group social dynamics. The platform test can also be used to explore neural activity associated with dominance and social hierarchy. Ultimately, the platform test will enhance our understanding of social dominance behavior and the neural underpinnings of social hierarchy in both sexes, such that we can develop better therapies to treat impairments in socioemotional processing and expression of social behaviors, which are found in multiple psychiatric illnesses.

## Author Contributions

**Sarah Mott:** Conceptualization, Methodology, Investigation, Data curation, Formal Analysis, Writing – original draft, review, and editing. **Arushi Saha:** Data curation, Writing – review and editing. **Abir Mohsin:** Data curation, writing – review and editing. **Sarah Magee:** Data curation, writing – review and editing. **Shyenne Grady:** Data curation, writing – review and editing. **Mia Keller:** Data curation, writing – review and editing. **Melissa Herman:** Conceptualization, Methodology, Writing – original draft, review, and editing, funding acquisition.

## Supplemental

**Supplemental Figure 1.**
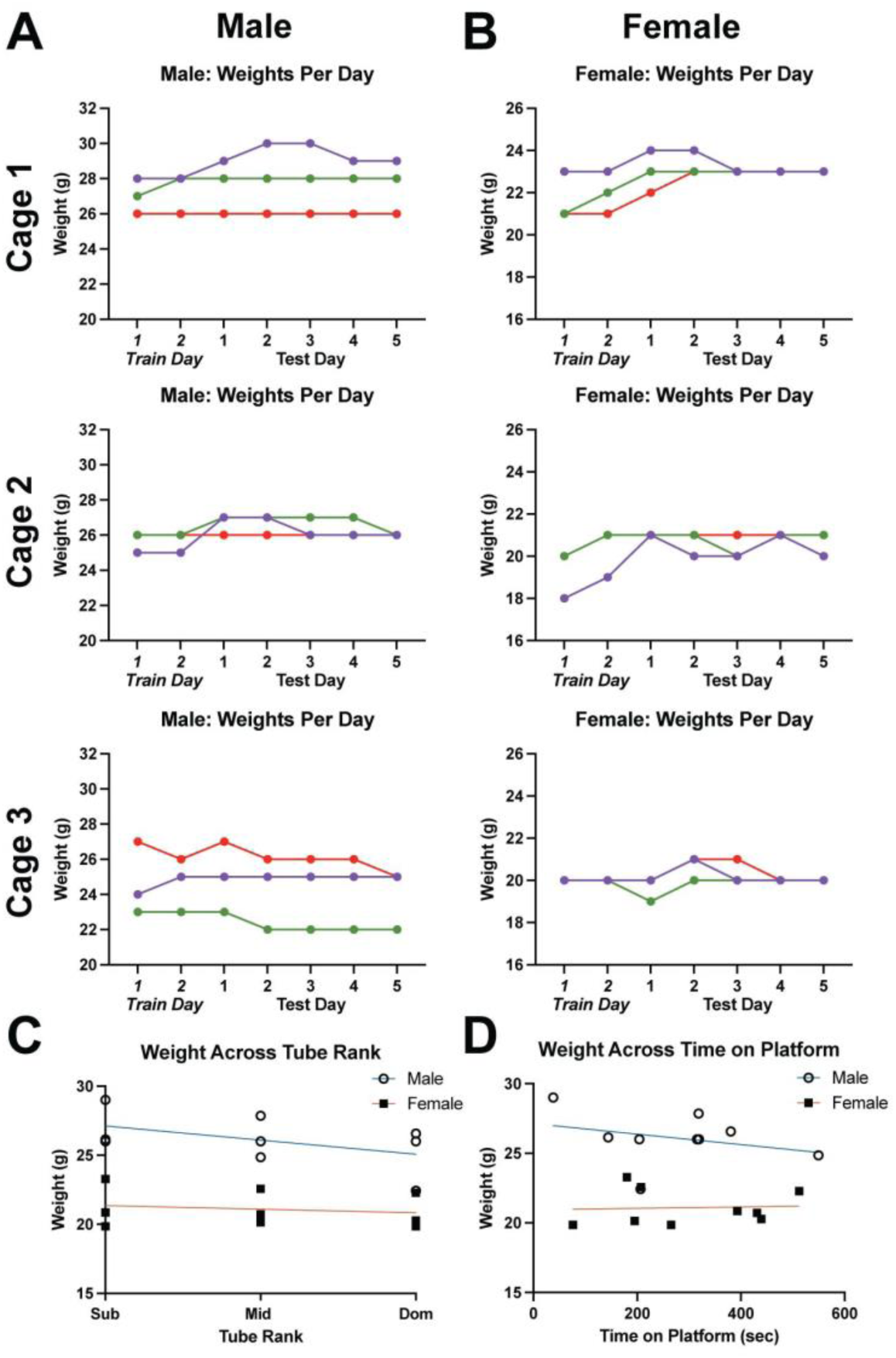
Weights. **A.** Daily weights of male cages. **B.** Daily weights of female cages. **C.** Regression analysis of average weight and tube rank for males and females (Simple linear regression: male: R^2^=0.2349, p=0.1861; female: R^2^=0.03112, p=0.6498). **D.** Regression analysis of average weight and time on platform for males and females (Simple linear regression: male: R^2^=0.09233, p=0.4267; female: R^2^=0.003380, p=0.8819).

**Supplemental Figure 2.**
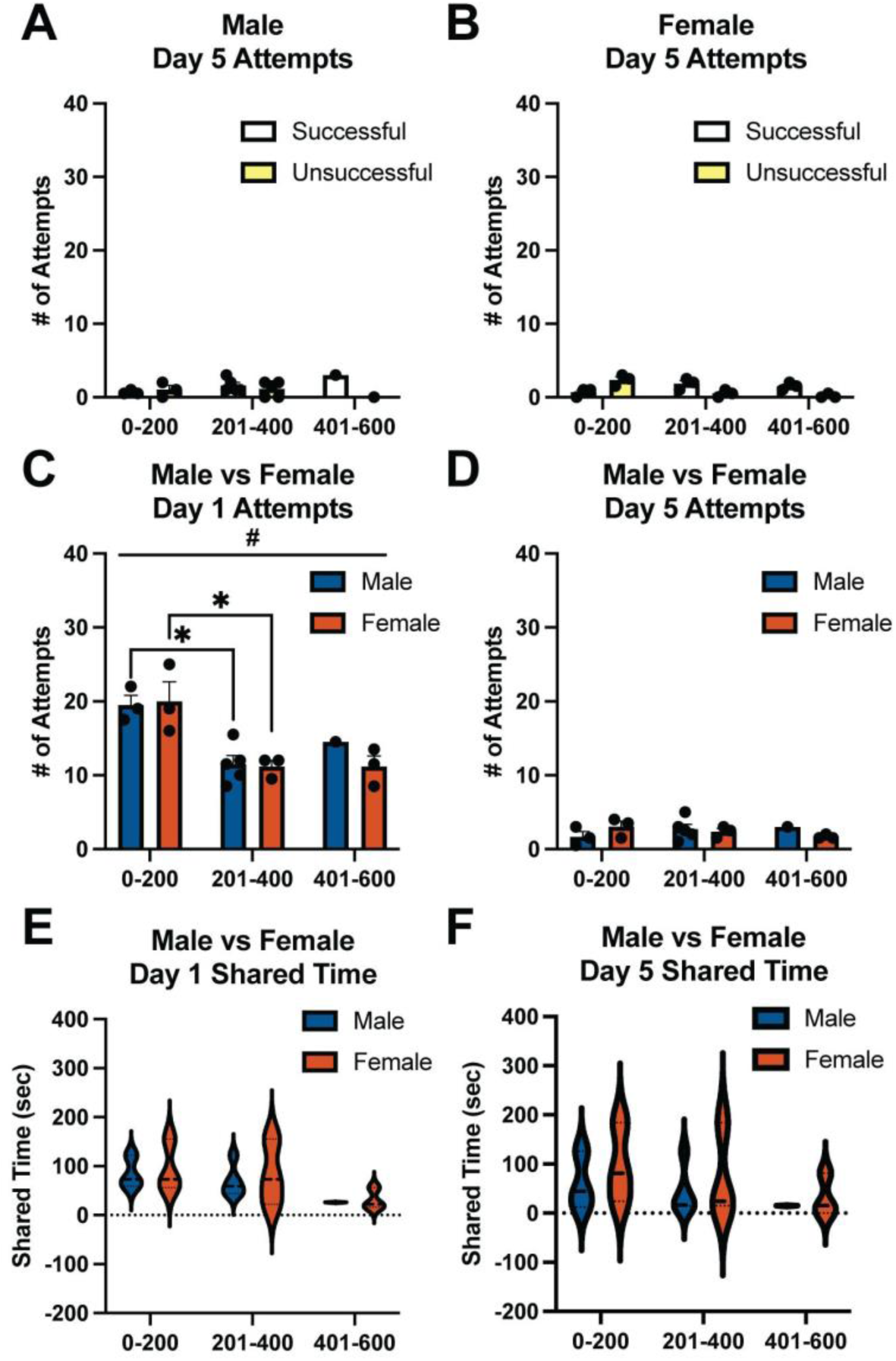
Motivated Behavior on Day 5 and Sex Differences. **A.** Types of attempts made by male mice to get on the platform on Day 5. **B.** Types of attempts made by female mice to get on the platform on Day 5. **C.** Total attempts made by males and females on Day 1 (2-way ANOVA: main effect of platform time: F(2,12)=15.40, #p=0.0005; Tukey’s, Male 0-200 v 201-400, p=0.0216; Tukey’s, Female 0-200 v 201-400, p=0.0224). **D.** Total attempts made by males and females on Day 5. **E.** Total time spent sharing the platform on Day 1 for males and females. **F.** Total time spent sharing the platform on Day 5 for males and females.

**Supplemental Figure 3.**
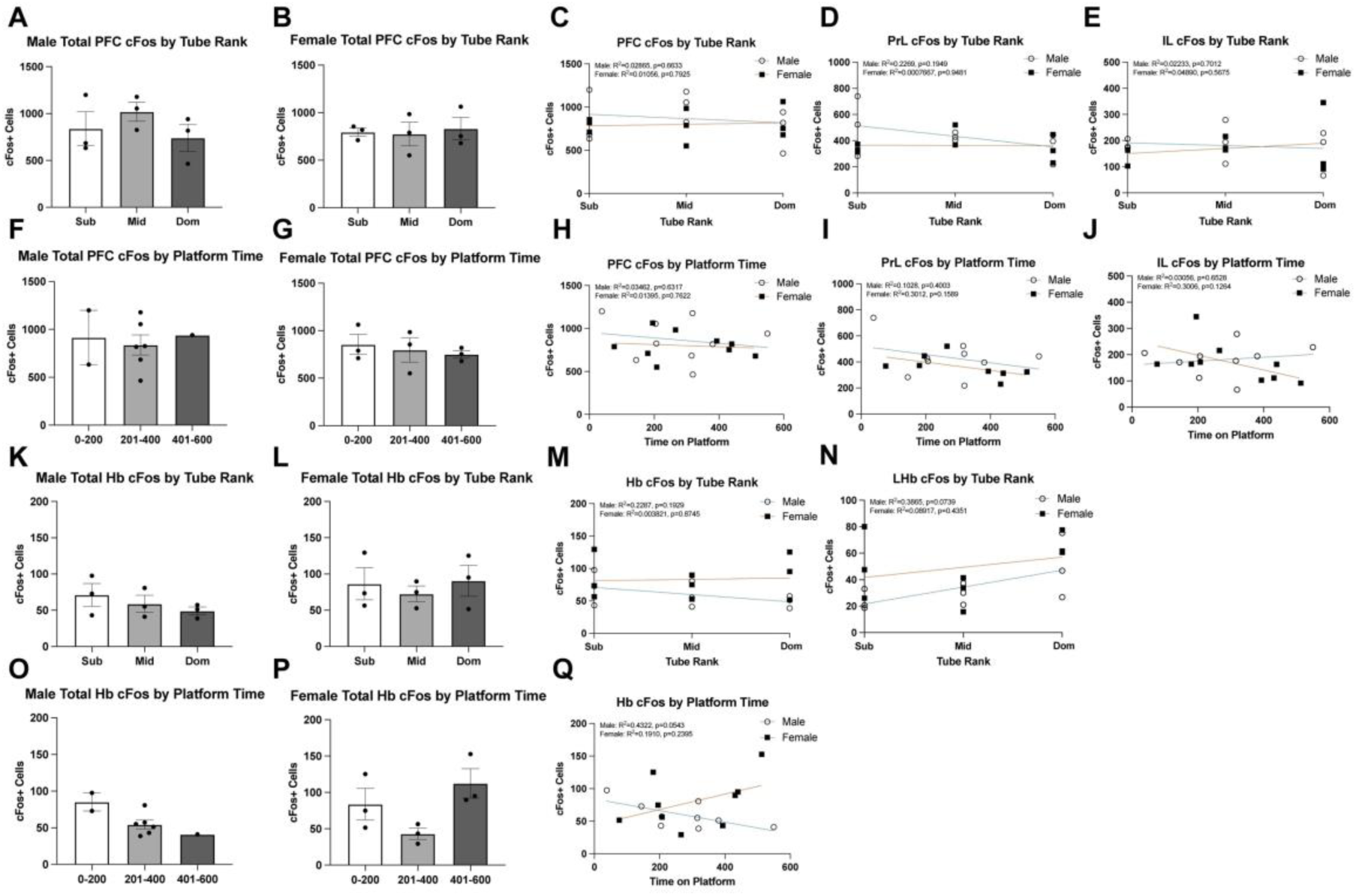
Total Neuronal Activity in PFC and Hb Following Tube and Platform. **A.** Total male PFC cFos+ cells averaged by animal for tube rankings. **B.** Total female PFC cFos+ cells averaged by animal for tube rankings. **C.** Regression analysis of total PFC cFos expression and tube rank for males and females (Simple linear regression: male: R^2^=0.02865, p=0.6633; female: R^2^=0.01056, p=0.7925). **D.** Regression analysis of cFos expression in the PrL subregion of the PFC and tube rank for males and females (Simple linear regression: male: R^2^=0.2269, p=0.1949; female: R^2^=0.0007667, p=0.9481). **E.** Regression analysis of cFos expression in the IL subregion of the PFC and tube rank for males and females (Simple linear regression: male: R^2^=0.02233, p=0.7012; female: R^2^=0.04890, p=0.5675). **F.** Total male PFC cFos+ cells averaged by animal for time on platform. **G.** Total female PFC cFos+ cells averaged by animal for time on platform. **H.** Regression analysis of total PFC cFos expression and time on platform for males and females (Simple linear regression: male: R^2^=0.03462, p=0.6317; female: R^2^=0.01395, p=0.7622). **I.** Regression analysis of cFos expression in the PrL subregion of the PFC and time on platform for males and females (Simple linear regression: male: R^2^=0.1028, p=0.4003; female: R^2^=0.3012, p=0.1589). **J.** Regression analysis of cFos expression in the IL subregion of the PFC and time on platform for males and females (Simple linear regression: male: R^2^=0.03056, p=0.6528; female: R^2^=0.3006, p=0.1264). **K.** Total male Hb cFos+ cells averaged by animal for tube rankings. **L.** Total female Hb cFos+ cells averaged by animal for tube rankings. **M.** Regression analysis of total Hb cFos expression and tube rank for males and females (Simple linear regression: male: R^2^=0.2287, p=0.1929; female: R^2^=0.003821, p=0.8745). **N.** Regression analysis of cFos expression in the LHb subregion and tube rank for males and females (Simple linear regression: male: R^2^=0.3865, p=0.0739; female: R^2^=0.08917, p=0.4351). **O.** Total male Hb cFos+ cells averaged by animal for time on platform. **P.** Total female Hb cFos+ cells averaged by animal for time on platform. **Q.** Regression analysis of total Hb cFos expression and time on platform for males and females (Simple linear regression: male: R^2^=0.4322, p=0.0543; female: R^2^=0.1910, p=0.2395).

**Supplemental Figure 4.**
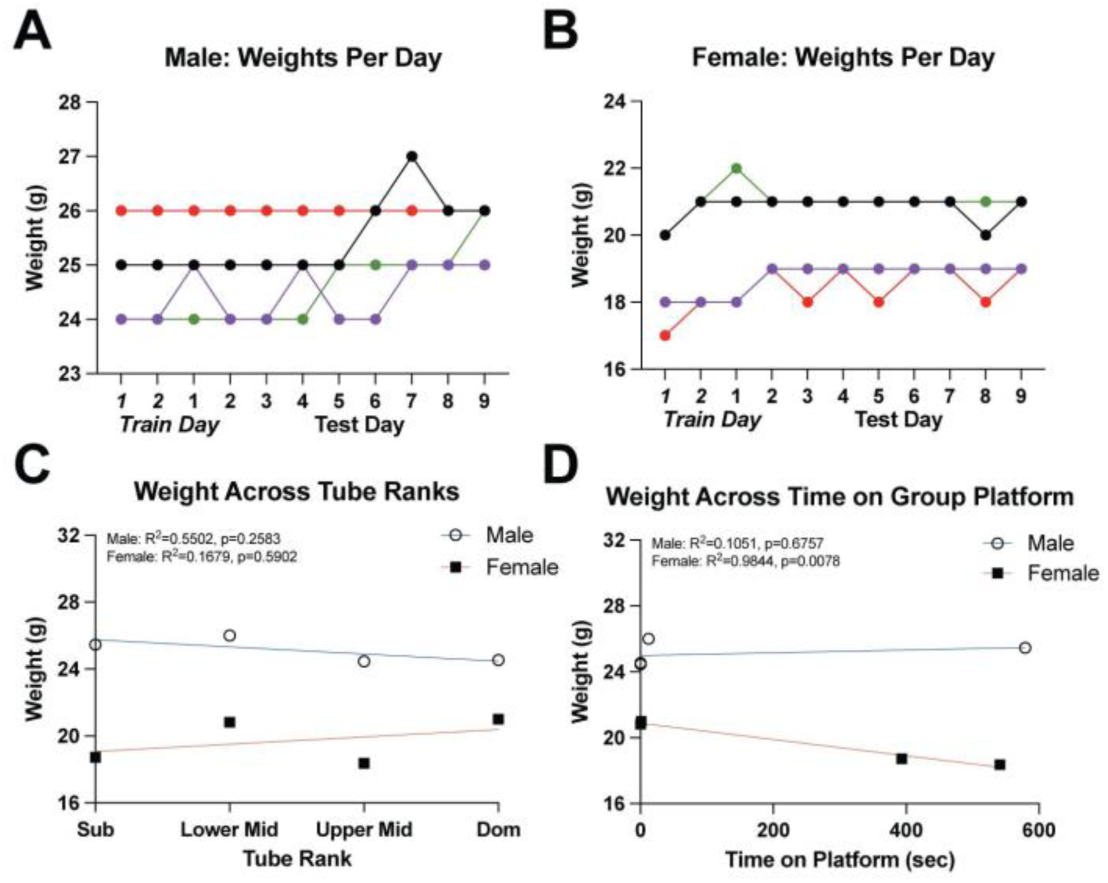
Group Platform Weights. **A.** Daily weights of male cage. **B.** Daily weights of female cage. **C.** Regression analysis of average weight and tube rank for males and females (Simple linear regression: male: R^2^=0.5502, p=0.2583; female: R^2^=0.1679, p=0.5902). **D.** Regression analysis of average weight and time on group platform for males and females (Simple linear regression: male: R^2^=0.1051, p=0.6757; female: R^2^=0.9844, p=0.0078).

**Supplemental Figure 5.**
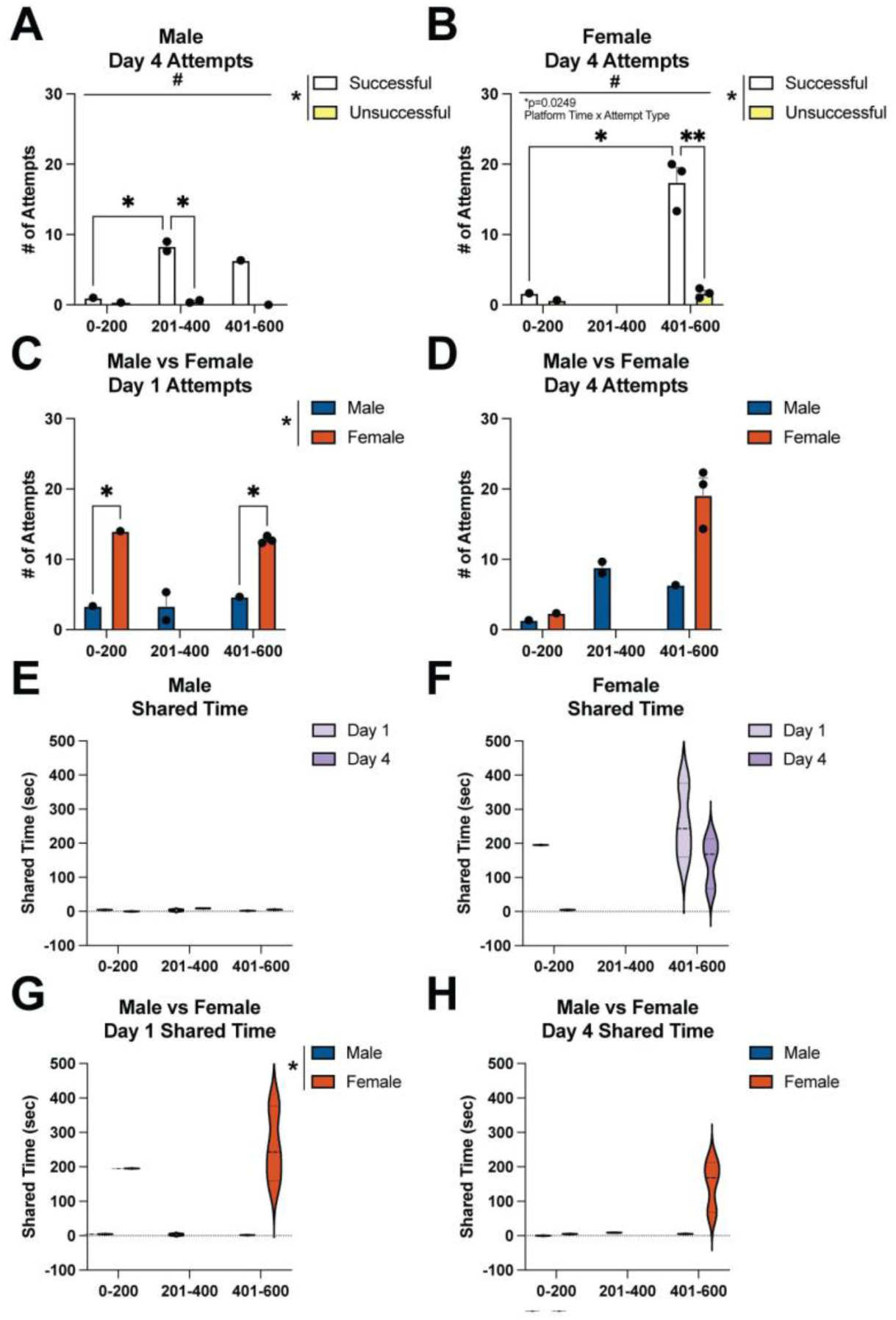
Paired Platform Motivated Behavior, Cooperative Behavior, and Sex Differences. **A.** Types of attempts by male mice to get on the platform on Day 4 (2-way ANOVA: main effect of platform time: F(2,2)=19.85, #p=0.0480; main effect of attempt type: F(1,2)=93.19, *p=0.0106; Tukey’s, Successful 0-200 v 201-400, p=0.0455; Tukey’s, 201-400 Successful v Unsuccessful, p=0.0269). **B.** Types of attempts made by female mice to get on the platform on Day 4 (2-way ANOVA: main effect of platform time: F(1,4)=15.79, #p=0.0165; main effect of attempt type: F(1,4)=15.79, *p=0.0165; significant interaction: F(1,4)=12.25, p=0.0249; Tukey’s, Successful 0-200 v 401-600, p=0.0208; Tukey’s, 401-600 Successful v Unsuccessful, p=0.0059). **C.** Total attempts made by males and females on Day 1 (2-way ANOVA: main effect of sex: F(1,4)=39.81, *p=0.0032; Tukey’s, 0-200 Male v Female, p=0.0186; Tukey’s, 401-600 Male v Female, p=0.0186). **D.** Total attempts made by males and females on Day 4. **E.** Total time spent sharing across paired platform testing in male mice. **F.** Total time spent sharing across paired platform testing in female mice. **G.** Total time spent sharing the platform on Day 1 for males and females (2-way ANOVA: main effect of sex: F(1,4)=10.46, *p=0.0319). **H.** Total time spent sharing the platform on Day 4 for males and females.

**Supplemental Figure 6.**
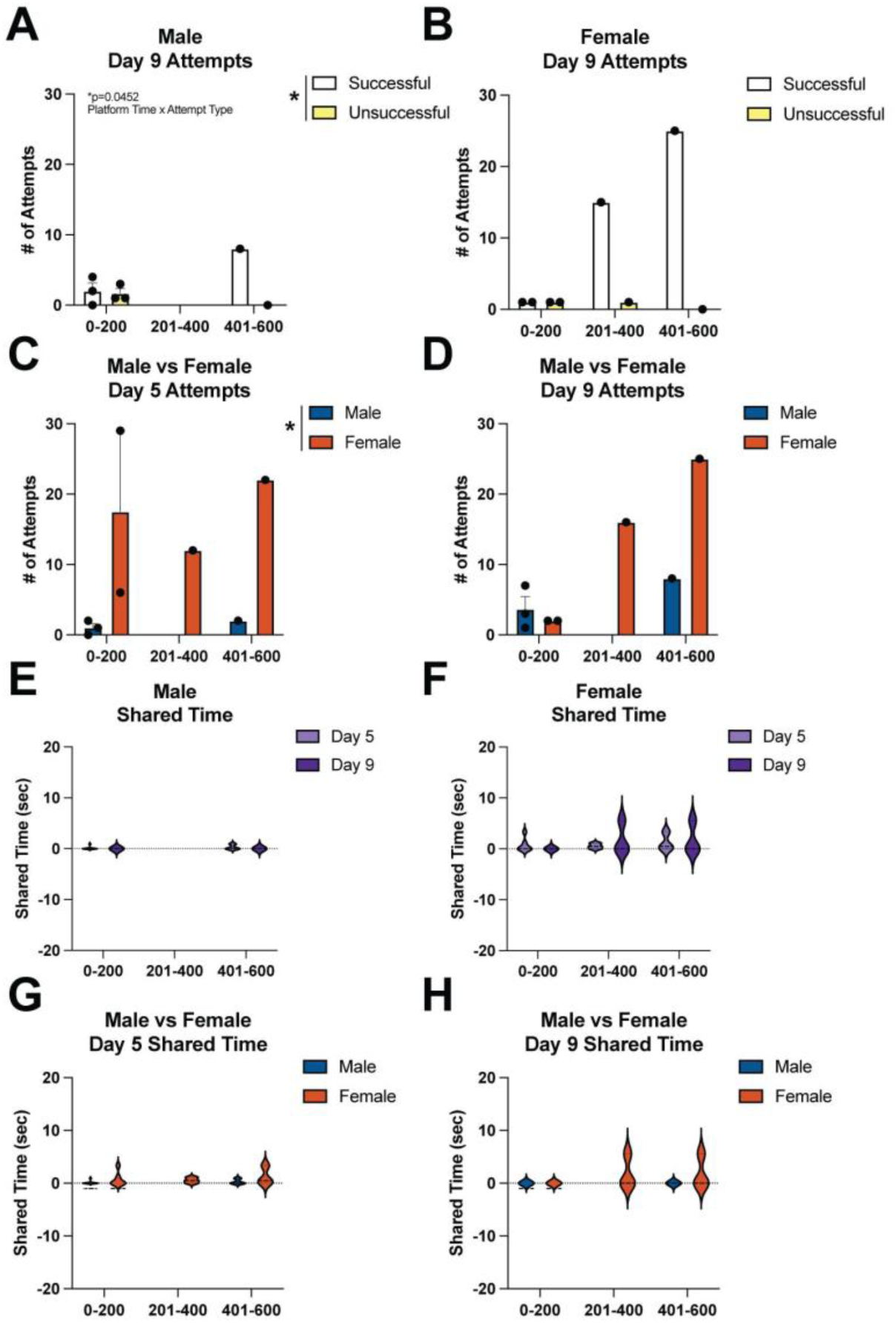
Group Platform Motivated Behavior, Cooperative Behavior, and Sex Differences. **A.** Types of attempts made by male mice to get on the platform on Day 9 (2-way ANOVA: main effect of attempt type: F(1,4)=9.766, *p=0.0354; significant interaction: F(1,4)=8.266, p=0.0452). **B.** Types of attempts made by female mice to get on the platform on Day 9. **C.** Total attempts made by males and females on Day 5 (2-way ANOVA: main effect of sex: F(1,4)=7.715,*p=0.0499). **D.** Total attempts made by males and females on Day 9. **E.** Total time spent sharing across group platform testing in male mice. **F.** Total time spent sharing across group platform testing in female mice. **G.** Total time spent sharing on Day 5 for males and females. **H.** Total time spent sharing the platform on Day 9 for males and females.

